# Sam50-Mic19-Mic60 axis Determines Mitochondrial Cristae Architecture by Mediating Mitochondrial Outer and Inner Membrane Contact

**DOI:** 10.1101/345959

**Authors:** Junhui Tang, Kuan Zhang, Jun Dong, Chaojun Yan, Shi Chen, Zhiyin Song

## Abstract

Mitochondrial cristae are critical for efficient oxidative phosphorylation, however, how cristae architecture is precisely organized remains largely unknown. Here, we discovered that Mic19, a core component of MICOS (mitochondrial contact site and cristae organizing system) complex, can be cleaved at N-terminal by mitochondrial protease OMA1. Mic19 directly interacts with mitochondrial outer-membrane protein Sam50 (the key subunit of SAM complex) and inner-membrane protein Mic60 (the key component of MICOS complex) to form Sam50-Mic19-Mic60 axis, which dominantly connects SAM and MICOS complexes to assemble MIB (mitochondrial intermembrane space bridging) supercomplex for mediating mitochondrial outer- and inner-membrane contact. OMA1-mediated Mic19 cleavage causes Sam50-Mic19-Mic60 axis disruption, which separates SAM and MICOS and leads to MIB disassembly. Disrupted Sam50-Mic19-Mic60 axis, even in the presence of SAM and MICOS complexes, causes the abnormal mitochondrial morphology, loss of mitochondrial cristae junctions, abnormal cristae distribution and reduced ATP production. Importantly, Sam50 displays punctate distribution at mitochondrial outer membrane, and acts as an anchoring point to guide the formation of mitochondrial cristae junctions. Therefore, we propose a model that Sam50-Mic19-Mic60 axis mediated SAM-MICOS complexes integration determines mitochondrial cristae architecture.

## INTRODUCTION

Mitochondria are fully articulated and highly organized organelles which surrounded by two membranes: the outer mitochondrial membrane (OMM) and the inner mitochondrial membrane (IMM) ^1^. The outer membrane is the first interface and barrier for mitochondria to communicate the substance, energy and information with the cytosol ^2^. The outer membrane possesses a dedicated protein import system, including TOM complex (translocase of the outer mitochondrial membrane), and SAM complex (mitochondrial sorting and assembly machinery) ^2, 3, 4^. The inner membrane is consists of the inner boundary membrane (IBM) and the cristae membrane (CM). OMM and IBM are firmly connected by contact sites. The IBM is closely apposed to the OMM while the inner membrane protrudes from the IBM into the inner space of the mitochondria formed cristae membrane ^5^, 6, 7. Cristae membranes are large tubular invaginations and are thought to increase the local charge density/pH to enhance ATP synthesis via oxidative phosphorylation ^8^. The connections between IBM and the cristae are the crista junctions (CJs), which are relatively uniform narrow, tubular and slot-like structures ^9^.

Cristae and crista junctions are important for mitochondrial organization and functions ^10^. Lots of experimental results revealed that the formation of cristae and crista junctions requires Mgm1 (known as OPA1 in mammals), the dimeric form of the F1FO-ATP synthase (F1FO), the MICOS (mitochondrial contact site and cristae organizing system) complex, and Prohibitins (PHBs). Mgm1, the dynamin-like fusion protein of IMM, mediates the fusion of IMM, it cooperates with dimeric F1FO to stabilize the cristae membranes and to thereby generate the sac-like structure ^9^, 11, 12. Assembly of the MICOS complex is proposed to limit the fusion process by forming a crista junction ^8, 13, 14^. Prohibitins exist as two closely related proteins (PHB1 and PHB2) that localize to the IMM ^15^. PHBs were reported to assemble into ring-like structures that provide a frame-work to stabilize the structure of crista membrane ^14, 16^. However, So far, how mitochondrial cristae and crista junctions are formed remains largely unknown.

Eight subunits of MICOS complex in mammalian are described: Mic60/IMMT, Mic19/CHCHD3, Mic10/MINOS1, Mic23/Mic26/APOO, Mic27/APOOL, Mic13/QIL1, Mic25/CHCHD6 and Mic14/CHCHD10 ^17, 18, 19, 20, 21, 22, 23, 24, 25^. Mic19 and Mic25 are peripheral membrane protein whereas other MICOS subunits are transmembrane proteins containing at least one transmembrane domain. In addition to the 700kDa integrated MICOS complex, some core components like Mic60, Mic19 and Mic25 can also from a proximately 450kDa subcomplex, which probably is the nucleation of MICOS assembling ^6, 26^. In MICOS complex, Mic60, Mic19 and Mic10 play a dominant role in cristae patterning ^17, 27^. Furthermore, MICOS subunits also possess other capacity: Mic10 and Mic60 show the ability in bending the liposome membrane in vitro ^28, 29^; Mic60 or Mic19 ablation can affect the mitochondrial dynamics ^17, 18, 30^; MICOS complex mediates the formation of mitochondrial contact site which is response for mitochondrial outer- and inner-membrane contact ^31^. In addition, Mic60, Mic19 and Mic25, which are responsible for mitochondrial contact sites formation, have the potential to bind the outer membrane protein Sam50 ^32, 33, 34^. Sam50 depletion could trigger abnormal mitochondrial morphology and cristae structures ^35, 36, 37^, but the mechanism of Sam50 in regulating cristae organization is still unknown. In addition, it remains unclear how the synergistic interactions between SAM and MICOS complex play role in the formation of mitochondrial contact sites and the biogenesis of mitochondrial cristae structure.

Recently, some labs reported a larger protein complex, around 2200–2800 kDa, stretching across the inner and out membrane which termed ‘mitochondrial intermembrane space bridging’ (MIB) complex ^26, 36, 38^, in which MICOS complex interacts with DNAJC11 and outer membrane proteins SAM50, Metaxin 1 (MTX1), Metaxin2 (MTX2) and Metaxin 3 (MTX3) to regulate the cristae function ^6, 39, 40^. However, the mechanism of MIB complex assembly and the role of MIB complex in mitochondrial morphology and structure are largely unknown.

In this study, we find that Mic19 can be cleaved at N-terminal by mitochondrial metalloprotease OMA1 after Sam50 depletion or CCCP treatment. This feature provides us with excellent tools to prove the key role of the Sam50-Mic19-Mic60 axis in mediating mitochondrial outer- and inner-membrane contact. The cleaved short form of Mic19 (S-Mic19) disrupts the interaction between SAM and MICOS and then causes the abnormal mitochondrial morphology, loss of mitochondrial crista junctions, and impairs ATP production. We also put forward the notion that Sam50 may serve as an anchoring point in OMM for the formation of mitochondrial cristae junction. Moreover, Mic25, an important paralog of Mic19, can act as reserves to rescue the interaction between Sam50 and Mic60. When Mic25 was overexpressed in Mic19 knockout cells, mitochondrial morphology and crista junctions can be reconstructed. Overall, we provide a new mode that Sam50-Mic19-Mic60 axis-mediated OMM and IMM contact dominantly regulates the formation of cristae junction.

## RESULTS

### Mic19 could be cleaved by mitochondrial protease OMA1

To maintain the mitochondrial homeostasis, there is a well-established enzymes system in this semi-autonomous organelle. Our previous works have shown that Mic60 degradation, caused by down-regulation of Mic19, is mediated by the mitochondrial protease Yme1L ^17^. Because Mic19 depletion also leads to the degradation of Sam50 ^18^, we test whether Yme1L mediates Sam50 degradation. Yme1L knockdown partly inhibited Sam50 degradation induced by Mic19 knockdown (Figure S1A), furthermore, Yme1L physically interacted with Sam50 (Figure S1B). These data suggest that Yme1L regulates Sam50 degradation. Because Sam50 directly binds to Mic19 ^36^, we examine whether Sam50 is related to Mic19 degradation. To our surprise, instead of causing degradation of Mic19, Sam50 knockdown (shSam50) resulted in the cleavage of Mic19 and formed a short form of Mic19 (S-Mic19) (Figure 1A). However, the depletion of Mic60 or Mic25 did not induce Mic19 cleavage (Figures S1C and S1D). Next, considering that Mic19 is a mitochondrial intermembrane space protein, we investigated whether mitochondrial inner membrane protease Yme1L or OMA1 is responsible for the cleavage of Mic19. Mic19 was cleaved in Yme1L knockout but not in OMA1 knockout cells (Figures 1A and 1B), suggesting that OMA1 regulates Mic19 cleavage.

**Figure 1.**
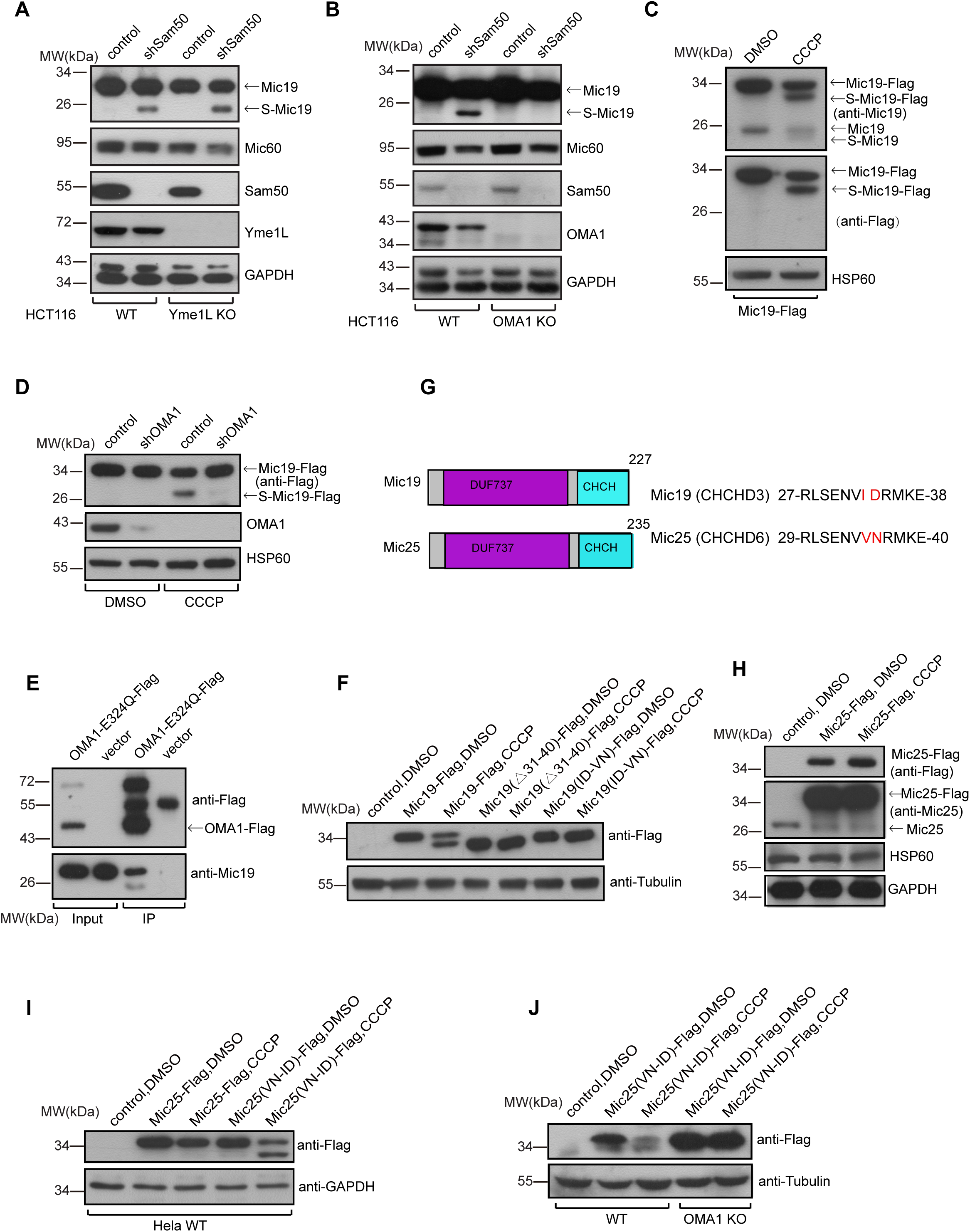
Mic19 could be cleaved by mitochondrial protease OMA1. (A) Sam50 knockdown (shSam50, 7 days) was performed in WT or Yme1L KO HCT116 cells respectively, and cell lysates were analyzed for the indicated protein expression by Western blotting. GAPDH was used as loading control. (B) Sam50 was depleted (shSam50, 7 days) in WT or OMA1 KO HCT116 cells respectively. Cell lysates were analyzed by Western blotting using the indicated antibodies. (C) HeLa cells stably expressing human Mic19-Flag were treated with DMSO or CCCP (20μM, 4 h), and the cell lysates were analyzed by Western blotting using ani-Mic19, anti-Flag or anti-HSP60 antibodies. (D) WT or OMA1 KO MEFs cells expressing Mic19-Flag were treated with DMSO or CCCP (20μM, 4 h). Mic19-Flag cleavage was examined by Western blotting using indicated antibodies. (E) Cell lysates of 293T cells expressing control or OMA1^E324Q^-Flag were Immunoprecipitated with anti-Flag M2 resin, and the eluted protein samples were analyzed by Western blotting using indicated antibodies. (F) HeLa cells expressing Mic19-Flag and indicated Mic19 mutants (Flag-tagged) were treated with DMSO or CCCP (20μM, 4 h), and the cell lysates were analyzed by Western blotting using anti-Flag or anti-Tubulin antibodies. “Δ” indicates deletion of residues. (G) The schematic presentation of human Mic19 and Mic25 domain. ‘DUF737’ indicates domain of unknown function; ‘CHCH’ indicates the coiled-coil helix-coiled-coil helix domain. The comparison of Mic19 N-terminal 27-38AA and Mic25 N-terminal 29-40AA were showed. (H-J) HeLa cells expressing Mic25-Flag (H) or indicated Mic25 mutants (I), and WT or OMA1 KO HCT116 cells expressing Mic25 mutants (J) were treated with DMSO or CCCP (20μM, 4 h). Cell lysates were then analyzed by Western blotting using the indicated antibodies.

To determine whether the Mic19 cleavage locates at the N-terminus or C-terminus, Mic19-Flag (Flag tag at the C-terminus) was transiently overexpressed in 293T cells. Unexpectedly, the cleavage of Mic19-Flag was also detected (Figure S1E). Due to little endogenous Mic19 cleavage in wild-type cells, we speculate that excessive Mic19-Flag, caused by transient expression, may not be completely assembled to MICOS complex, and leaving unassembled Mic19 or Mic19-Flag is then cleaved by OMA1. To examine our hypothesis, we built the Mic19-Flag stably overexpressed HeLa cells and treated cells with CCCP (an oxidative phosphorylation inhibitor, capable of reducing mitochondrial membrane potential and activating OMA1 metalloprotease activity ^41^. As we expected, both endogenous S-Mic19 and exogenous S-Mic19-Flag were detected in CCCP-treatment group (Figure 1C), implying unassembled Mic19 can be cleaved at N-terminal. In contrast, in response to OMA1 knockdown (in HeLa cells) or OMA1 knockout (in MEFs), overexpressed Mic19-Flag failed to be cleaved even after CCCP treatment (Figures 1D and S1F). In addition, co-immunoprecipitation assay revealed that Mic19 interacted with OMA1^E324Q^, a mutation that only blocks the OMA1 protease activity ^42^ (Figure 1E). These results demonstrated that Mic19 is cleaved at N-terminal by mitochondrial protease OMA1 in response to Sam50 downregulation or CCCP treatment.

To ascertain the cutting site of Mic19, we first compared the molecular weight of proteins being displayed in SDS-PAGE, and got an around 3kD size difference that is equivalent to 36 amino acids on average. Furthermore, alignment of amino acid sequence at N terminal of Mic19 from different species showed that the region (27–38aa) of homo Mic19 is highly evolutionarily conserved (Figure S1G). We then deleted 31-40aa of Mic19 and found the N-terminal processing of Mic19 is inhibited after CCCP treatment (Figure 1F). Interestingly, Mic25, another MICOS subunit, is about 90 percent homologous with the region (27–38aa) of Mic19, and only two amino acids ‘VN’ of Mic25 is different from that in Mic19 (Figure 1G). Meanwhile, we searched the downstream sequence right after the site number 39 and found that Mic19 protein sequence in some species (like Equus caballus) starts from site 36 of human Mic19 right behind the two un-conserved amino acids (Figure S1G). The un-conserved character within this sequence indicates that these two amino acids may be critical for cleavage of Mic19. Importantly, exogenous Mic25-Flag is un-cleavable upon CCCP treatment (Figure 1H), so we mutated ‘ID’ in Mic19 to ‘VN’ (Mic19^ID^33-34^VN^) and implemented in a manner similar as the control did (Figure 1G). No cleavage at the N-terminal of Mic19^ID^33-34^VN^ was detected in response to CCCP treatment (Figure 1G), indicating that amino acids ‘ID’ of Mic19 is the site of cleavage. To eliminate the possibility that this two amino acids are important for OMA1 binding but not cleavage, we co-expressed Mic19-Flag or Mic19^ID^33-34^VN^-Flag with OMA1^E324Q^-Myc in 293T cells, co-immunoprecipitation assay and followed immunoblotting revealed that both Mic19-Flag and Mic19^ID^33-34^VN^-Flag interacted with OMA1^E324Q^-Myc (Figures S1H), demonstrating that ‘ID’ is OMA1 cleavage site but not binding site. To further test whether two amino acids are indispensable for the OMA1 cleavage, we replaced the ‘VN’ in Mic25 with ‘ID’ (Mic25^VN^35-36^ID^) and examined the processing of Mic25^VN^35-36^ID^. OMA1 dependent cleavage of Mic25^VN^35-36^ID^ was displayed (Figures 1I and 1J). Therefore, the inter-paralog replacement of two amino acids endows Mic19 obvious resistance and Mic25 more susceptible to OMA1 cleavage. Overall, the processing site of Mic19 by OMA1 is possibly right after the ‘ID’ at its N terminal.

### Mic19 cleavage disrupts Sam50-Mic19-Mic60 axis and disassembles MIB supercomplex

As an important subunit of the MICOS complex, previous work has shown that Mic19 interacts with the mitochondrial outer membrane Sam50 and the mitochondrial inner membrane Mic60 ^18^. We also demonstrated the interaction again by mass spectrometry, co-immunoprecipitation (co-IP) assay and GST-pull down assay (Supplementary Table S1 and Figures S2A-2C). Therefore, we firstly investigated whether Mic19 cleavage affects the interactions between Mic19 and other proteins. Co-IP assay revealed that Mic60 interacts with Mic19-Flag and S-Mic19-Flag but not Mic19 (1-35aa)-Flag (N-terminal 35 amino acids of Mic19) (Figure 2A, lane 6, 7, 8). In contrast, Co-IP data showed that large amounts of Sam50 were precipitated by Mic19-Flag or Mic19 (1-35aa)-Flag, but little amount of Sam50 was precipitated by S-Mic19-Flag (Figure 2A, lane 6, 7, 8). A small amounts of Sam50 was still precipitated by S-Mic19-Flag probably because S-Mic19-Flag can interact with endogenous Mic19, which may combine this little amount of Sam50. Indeed, Mic19-Flag could precipitate S-Mic19 (Figure 2A, lane 6). These data suggest that Mic19 (1-35aa)-Flag but not S-Mic19-Flag interacts with Sam50. Next, GST-pull down assay verified that Mic19 (1-35aa)-Flag but not S-Mic19-Flag specifically binds to Sam50, and S-Mic19-Flag but not Mic19 (1-35aa)-Flag directly interacted with Mic60 (Figures 2B-2D). Therefore, these results demonstrated that the interaction between Mic19 and Sam50 was disrupted after Mic19 cleavage.

**Figure 2.**
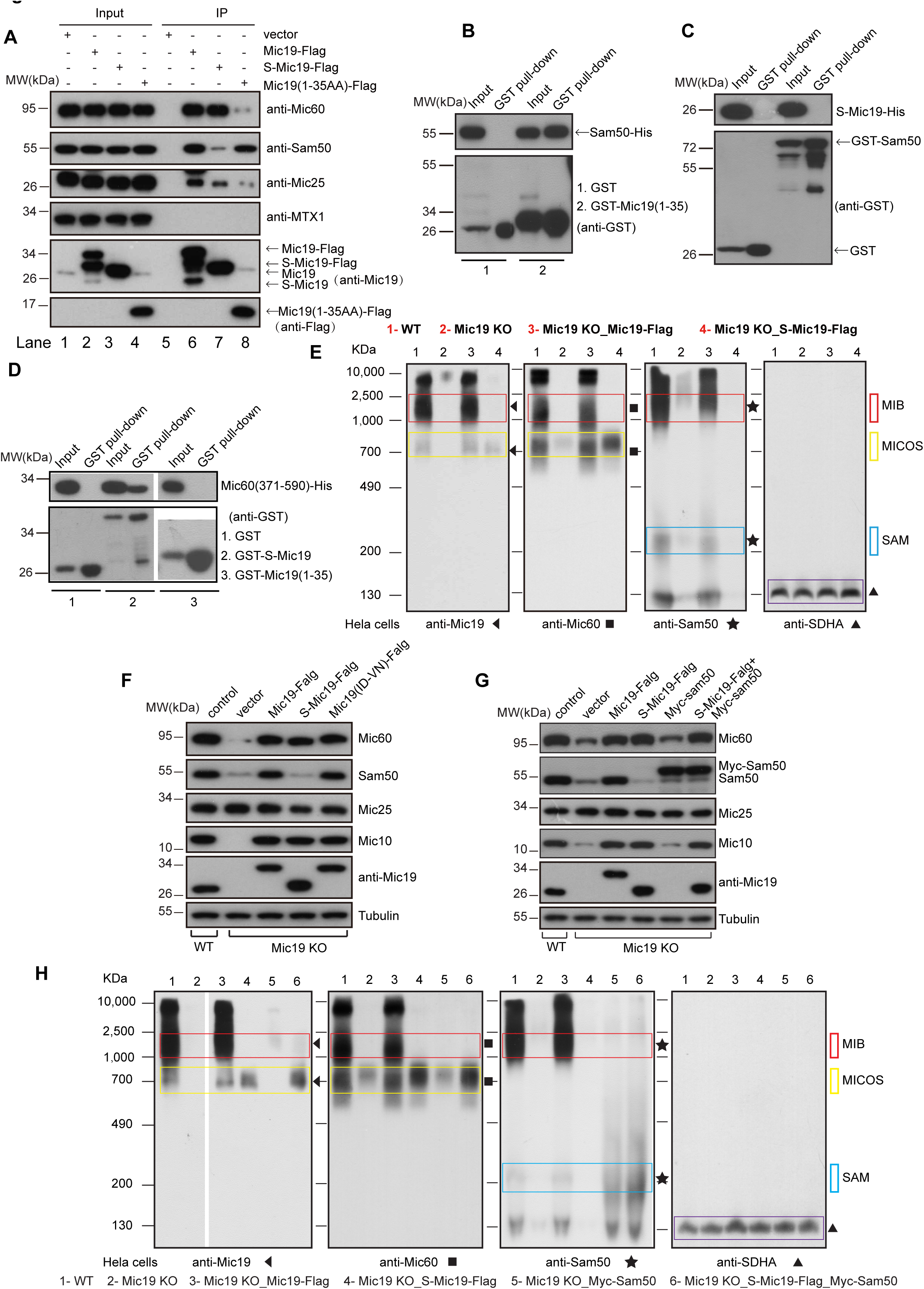
Mic19 cleavage disrupts Sam50-Mic19-Mic60 axis and disassembles the MIB supercomplex. (A) Cell lysates of 293T cells expressing Mic19-Flag, S-Mic19-Flag, and Mic19 (1-35aa)-Flag were used for co-immunoprecipitation (co-IP) assay with anti-Flag M2 resin, and the protein samples were subjected to immunoblot using the indicated antibodies. (B-D) GST or His fused proteins were expressed in E. coli, cell extracts were used for GST pull-down assay by using the Pierce Glutathione Agarose, and the protein samples were analyzed by Western blotting using antibodies against GST or His. The examination of the direct interaction between Mic19 (1-35AA) and Sam50 (B). The demonstration of the direct interaction between S-Mic19 and Sam50 (C). The detection of direct interaction between Mic19 (1-35AA) or S-Mic19 with Mic60 (371-590AA) (D). (E) Mitochondria were extracted from WT, Mic19 KO, or Mic19^ΔgRNA^-Flag or S-Mic19^ΔgRNA^-Flag expressed Mic19 KO HeLa cells, then non-denatured protein samples from mitochondria were analyzed for MIB, MICOS and SAM complexes by BN-PAGE and Western blotting using indicated antibodies. The bands of the complexes are labeled with corresponding boxes. The MIB complex is approximately 2000kDa, and the MICOS is approximately 700kDa. SAM complex is approximately 250kDa, ∼125kDa unidentified complex containing Sam50 is showed. Mitochondrial supercomplex II is detected by anti-SDHA, served as loading control. (ΔgRNA indicates mutation in gRNA targeting site, without altering the amino acid sequences). (F) Mic19^ΔgRNA^-Flag, S-Mic19^ΔgRNA^-Flag, and Mic19^(ID^33-34^VN)ΔgRNA^-Flag were stably expressed in Mic19 KO HeLa cells respectively. Cell lysates were analyzed by Western blot using indicated antibodies. (G) Myc-Sam50 were stably expressed in Mic19 KO, or S-Mic19^ΔgRNA^-Flag expressed Mic19 KO HeLa cells, the cell lysates were analyzed for expression of MICOS subunits by Western blotting using indicated antibodies. (H) Mitochondria were extracted from indicated cells, BN-PAGE and Western blotting analysis for the related cell lines, the non-denatured protein samples from mitochondria were analyzed for MIB, MICOS and SAM complexes by BN-PAGE and Western blotting. The bands of the complexes are labeled with corresponding boxes.

Sam50 is the core subunit of the SAM complex ^3^, and Mic60 is the key component of the MICOS complex ^13^, and it has been reported that there exists ‘mitochondrial intermembrane space bridging’(MIB) supercomplex mediated by the interaction of SAM and MICOS complexes by using complexome profiling analysis ^6^. We firstly performed BN-PAGE assay to confirm the SAM, MICOS and MIB complexes by using Mic19 knockout HeLa cells and the Mic19 cardiac-specific knockout mice heart (Figures S2D and S2E). Next, we examined the effect of Mic19 cleavage on the assembly of these complexes. Mic19^ΔgRNA^-Flag and S-Mic19^ΔgRNA^-Flag were reintroduced in Mic19 KO HeLa cells respectively, and mitochondria from WT cells, Mic19 KO cells, Mic19^ΔgRNA^-Flag cells and S-Mic19^ΔgRNA^-Flag expressed Mic19 KO cells were then analyzed by BN-PAGE. MIB complex was detected in WT (lane 1) or Mic19^ΔgRNA^-Flag expressed Mic19 KO cells (lane 3) but not in Mic19 KO (lane 2) or S-Mic19^ΔgRNA^-Flag expressed Mic19 KO cells (lane 4) (Figure 2E). Interestingly, MICOS complex (∼700kDa) were still maintained in S-Mic19^ΔgRNA^-Flag expressed Mic19 KO cells (Figure 2E), suggesting that S-Mic19 is enough for assembly of MICOS complex. In addition, immunoblotting assay revealed that MICOS subunits Mic60 and Mic10 were recovered in Mic19^ΔgRNA^-Flag, Mic19 ^(ID^33-34^VN) ΔgRNA^-Flag and S-Mic19^ΔgRNA^-Flag expressed Mic19 KO cells (Figure 2F). However, Sam50 was only recovered in Mic19^ΔgRNA^-Flag and Mic19^-(ID^33-34^VN)-ΔgRNA^-Flag but not in S-Mic19^ΔgRNA^-Flag expressed Mic19 KO cells (Figure 2F), suggesting that MIB supercomplex is disrupted in S-Mic19^ΔgRNA^-Flag expressed Mic19 KO cells because of the disruption of Sam50-Mic19 interaction. Therefore, Mic19 cleavage disrupts the integrity of MIB supercomplex. To further clarify that the interaction between Mic19 and Sam50 is required for the complex, we reintroduced Myc-Sam50 in S-Mic19^ΔgRNA^-Flag expressed Mic19 KO cells (Figure 2G). BN-PAGE assays revealed that S-Mic19^ΔgRNA^-Flag and Myc-Sam50 co-expressed Mic19 KO cells still loss MIB supercomplex although SAM and MICOS complexes are present (Figure 2H). Therefore, Mic19 cleavage disrupts the Sam50-Mic19-Mic60 axis and then result in the disassembly of MIB supercomplex.

It has been reported that Mic60 and Mic25 also bind to Sam50 ^32, 33, 34^, indicating that Sam50-Mic60 axis and Sam50-Mic25-Mic60 axis may also be responsible for MIB complex assembly. However, Sam50 is destabilized in S-Mic19^ΔgRNA^-Flag expressed Mic19 KO cells which maintains the normal level of Mic60 and Mic25 (Figures 2F and 2G), suggesting that Mic19-Sam50 but not Mic60-Sam50 or Mic25-Sam50 interaction is responsible for Sam50 stabilization. Moreover, MIB supercomplex was also disrupted even in the presence of SAM and MICOS complexes in S-Mic19^ΔgRNA^-Flag expressed Mic19 KO cells (Figure 2H), further confirmed that Sam50-Mic19-Mic60 axis dominantly contributes to MIB supercomplex assembly and mitochondrial outer- and inner-membrane contact.

### Disruption of Sam50-Mic19-Mic60 axis causes abnormal mitochondrial morphology and loss of crista junctions

Because Sam50-Mic19-Mic60 axis plays the dominant role in MIB supercomplex assembly and mitochondrial outer- and inner-membrane contact, we then investigated the role of this axis in mitochondrial morphology and structure. Mic19^ΔgRNA^-Flag, S-Mic19^ΔgRNA^-Flag or Mic19^-(ID^33-34^VN) ΔgRNA^-Flag was expressed in Mic19 KO HeLa cells. Mitochondrial morphology of cells stably expressing mitochondrial matrix-targeted GFP (mito-GFP) was visualized by confocal microscopy. Remarkably, almost all the Mic19 KO HeLa cells showed the ‘Expanded and spherical’, but not fragmented, mitochondrial network (Figure 3A). After re-expression of Mic19^ΔgRNA^-Flag or Mic19^-(ID^33-34^VN) ΔgRNA^-Flag in Mic19 KO cells, which restored Sam50-Mic19-Mic60 axis, the normal tubular mitochondria morphology was recovered (Figures 3A and 3B). Surprisingly, more ‘large spherical mitochondria’ appeared in S-Mic19^ΔgRNA^-Flag expressed-Mic19 KO cells in which Mic19-Mic60 interaction is restored but Sam50 is not recovered (Figures 2A, 2F, 3A, and 3B). We and Kozjak-Pavlovic lab have found that Sam50 depletion could cause abnormal spherical mitochondria ^36, 37^. To rule out the possibility of Sam50 deficiency causing abnormal mitochondrial morphology, and further confirm the Mic19-Sam50 interaction is required for the mitochondrial morphology, Myc-Sam50 was reintroduced in Mic19 KO cells or S-Mic19^ΔgRNA^-Flag expressed Mic19 KO cells. However, these cells expressing Myc-Sam50 still showed spherical mitochondria (Figures 3C and 3D). In addition, we previously reported that “large spherical” mitochondria were observed in Mic60 or Sam50 knockdown cells ^17, 37^. Therefore, our data demonstrate that the disruption of Sam50-Mic19-Mic60 axis results in abnormal spherical mitochondria.

**Figure 3.**
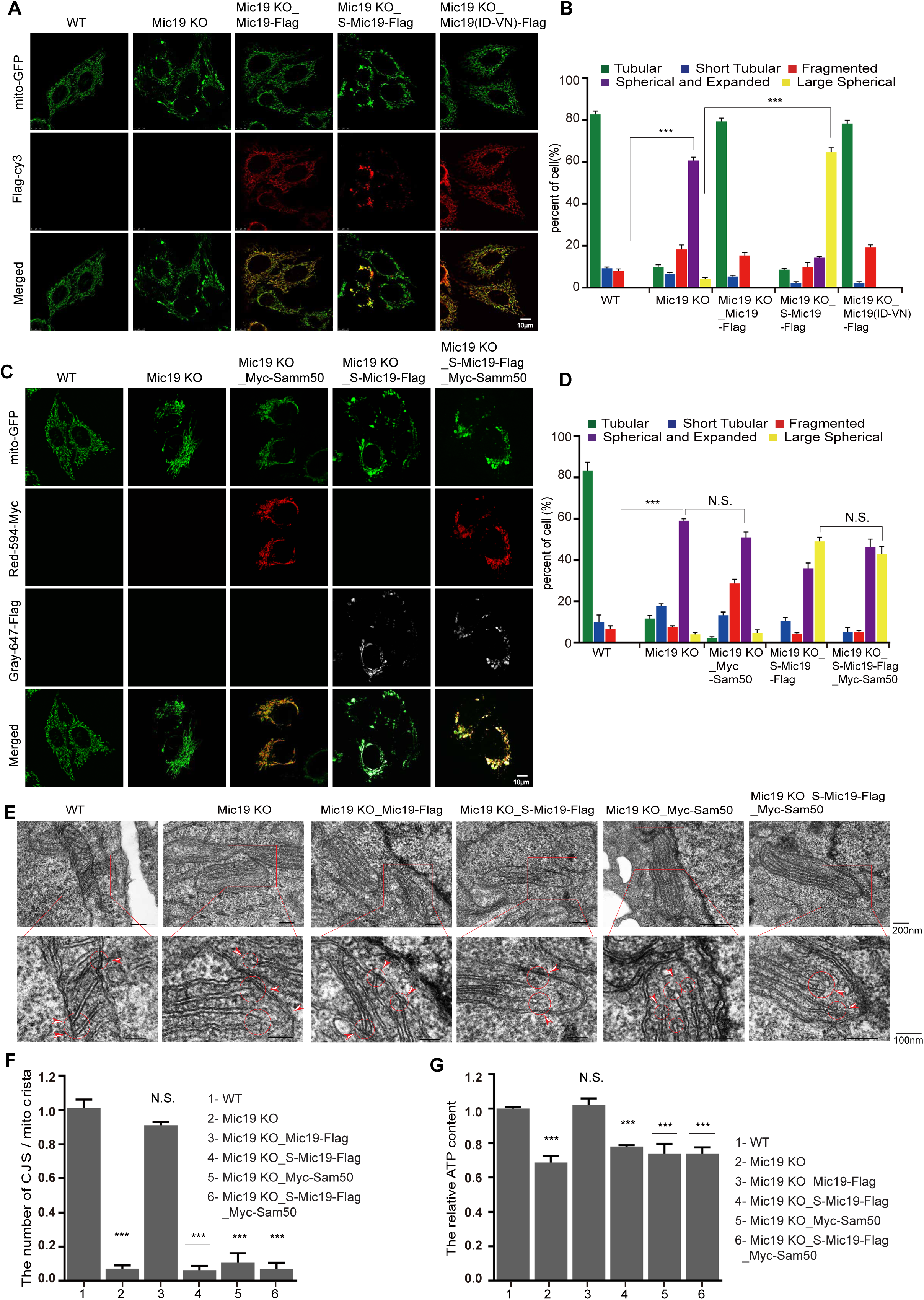
Disruption of Sam50-Mic19-Mic60 axis leads to abnormal mitochondrial morphology and loss of crista junctions. (A) Mic19 KO HeLa cells were expressed Mic19^ΔgRNA^-Flag, S-Mic19^ΔgRNA^-Flag or Mic19 ^(ID^33-34^VN)ΔgRNA^-Flag respectively, and mitochondrial morphology of all cell lines expressing mito-GFP (a mitochondrial marker) was visualized by confocal microscope. (B) Mitochondrial morphology described in (A) was counted according to the criteria detailed in “Materials and Methods”. Error bars indicate the mean ±SD of three independent experiments in which 100 cells were scored, ***P<0.001. (C) S-Mic19^ΔgRNA^-Flag, Myc-Sam50, or S-Mic19^ΔgRNA^-Flag plus Myc-Sam50 were expressed in Mic19 KO HeLa cells respectively. Mitochondrial morphology of indicated cells expressing mito-GFP was visualized by confocal microscope. (D) Mitochondrial morphology described in (C) was counted according to the criteria detailed in “Materials and Methods”. Statistical significance was assessed by Student’s t-test, error bars indicate the mean ±SD of three independent experiments in which 100 cells were scored, ***P<0.001. (E) Mitochondrial cristae junctions structure in WT, Mic19 KO, and Mic19 KO HeLa cells expressing Mic19-Flag, S-Mic19-Flag, Myc-Sam50 or S-Mic19-Flag plus Myc-Sam50 were analyzed by transmission electron microscope (TEM). The ultrathin section sample was observed using an electron microscope (JEM-1400plus, Tokyo, Japan) under the condition of an accelerating voltage of 80 kV. (F) Statistics of the mitochondrial ultrastructure in the cells described in (E). Error bars represent means ± SD of three independent experiments in which 100 mitochondrial cristae and the number of corresponding CJs were counted. ***P<0.001 vs WT. (G) The relative ATP level of WT, Mic19 KO, and Mic19 KO HeLa cells expressing Mic19-Flag, S-Mic19-Flag, Myc-Sam50 or S-Mic19-Flag plus Myc-Sam50 were measured using an ATP assay kit. Error bars represent means ± SD of three independent experiments, ***P<0.001 vs WT.

We and others have reported that deficiency of MICOS complex key subunits such as Mic60 or Mic10 affects mitochondrial cristae conformation ^17, 28^. Since Mic60 and Mic10 expression were recovered in S-Mic19^ΔgRNA^-Flag expressed Mic19 KO cells, we speculated that cristae should be normal. However, transmission electron microscopy (TEM) revealed that the crista junctions were still disappeared in S-Mic19^ΔgRNA^-Flag expressed Mic19 KO cells (Figures 3E). To further rule out the possibility that the decrease of Sam50 resulted in the disappearance of crista junctions (CJs), we next re-introduced Myc-Sam50 into S-Mic19^ΔgRNA^-Flag expressed Mic19 KO cells to directly confirm the role of the interaction between Mic19 and Sam50 in cristae organization. S-Mic19^ΔgRNA^-Flag and Myc-Sam50 co-expressed Mic19 KO cells maintained Sam50, Mic60 and Mic10 expression (Figure 2G), but still could not restore crista junctions (CJs) (Figures 3E and 3F). It is well known that mitochondrial energy production are closely related to mitochondrial cristae ^43^. We then measured ATP production in the cells. The disruption of the Sam50-Mic19 axis, even in the presence of SAM and MICOS complexes (Figure 2H), resulted in significantly reduced ATP production (Figure 3G). Therefore, Sam50-Mic19-Mic60 axis plays a critical role in the regulation of mitochondrial morphology, crista junctions (CJs) formation, and ATP production.

### Sam50 may act as a mitochondrial outer membrane anchoring point guiding the formation of crista junctions through Sam50-Mic19-Mic60 axis

Since Sam50-Mic19-Mic60 axis is critical for the maintenance of crista junctions, what is the role of Sam50, an outer membrane protein, in the formation of crista junctions? Previous works have reported that long-term Sam50 depletion leads to almost the complete loss of cristae ^36, 37^, probably due to indirect effects since TOM40 and VDACs are affected. Therefore, we assessed mitochondrial morphology and structure in short-term depletion of Sam50 (shSam50, 5 days) cells, which maintain the normal level of TOM40 and VDACs ^37^. In addition, to explore whether Mic19 cleavage induced by Sam50 deficiency is the primary effect leading to cristae deformation, we performed short-term Sam50 knockdown in Mic19^-(ID^33-34^VN) ΔgRNA^-Flag expressed Mic19 KO cells in which Mic19 cleavage is blocked. In response to Sam50 depletion (5 days), Mic60 and Mic10 were unaltered no matter whether Mic19 is cleaved or not (Figure 4A). Confocal microscopy revealed that shSam50 (5days) cells, shSam50 plus Mic19^ΔgRNA^-Flag or Mic19^-(ID^33-34^VN) ΔgRNA^-Flag expressed Mic19 KO cells displayed remarkably fragmented mitochondria (Figures 4B and 4C). These results suggest that Sam50 deficiency, but not the cleavage of Mic19, directly regulates mitochondrial morphology. Mitochondrial ultrastructure was also analyzed by transmission electron microscope (TEM). Most cristae of WT mitochondria connected with inner membrane and crista junctions were regularly arranged (Figure 4D). However, almost all cristae of shSam50 mitochondria pinched off from the inner membrane, leading to crista junctions collapsed (Figure 4E), the similar cristae were also found in shSam50 plus Mic19^ΔgRNA^-Flag or Mic19^-(ID^33-34^VN) ΔgRNA^-Flag expressed Mic19 KO cells (Figures 4E and 4F). These results indicate that Sam50 depletion is the primary effect leading to cristae deformation. Thus, we speculated that Sam50 might act as an anchoring point in mitochondria outer membrane to guide the formation of crista junctions. To further verify this hypothesis, we explored the localization of Sam50. Flag-Sam50 stably expressed COS7 cells were analyzed by using stimulated emission depletion microscopy (STED) techniques. The results showed that the distribution of Flag-Sam50 on mitochondrial outer membrane is dotted (Figure 4G). Importantly, statistical analysis showed that the location of about 30% Flag-Sam50 are adjacent to crista junctions (Figure 4H). Sastri et al. recently reported that about 16% of the crista junctions overlap with the Sam50-APEX2 positive domains in the inner membrane space ^34^. It is probably because Sam50 also acts as the translocase responsible for biogenesis of outer membrane β-barrel proteins. In addition, overexpressed Sam50-APEX2 (Sam50-APEX2 is about 80kDa, Flag-Sam50 is about 60kDa, and Sam50 is about 50kDa) might also influence the localization of Sam50 in mitochondrial outer membrane. Overall, Sam50 may serve as an outer membrane anchoring point for MICOS complex, and SAM connects MICOS through the sam50-Mic19-Mic60 axis to anchor the crista junctions at a specific site of mitochondrial inner membrane.

**Figure 4.**
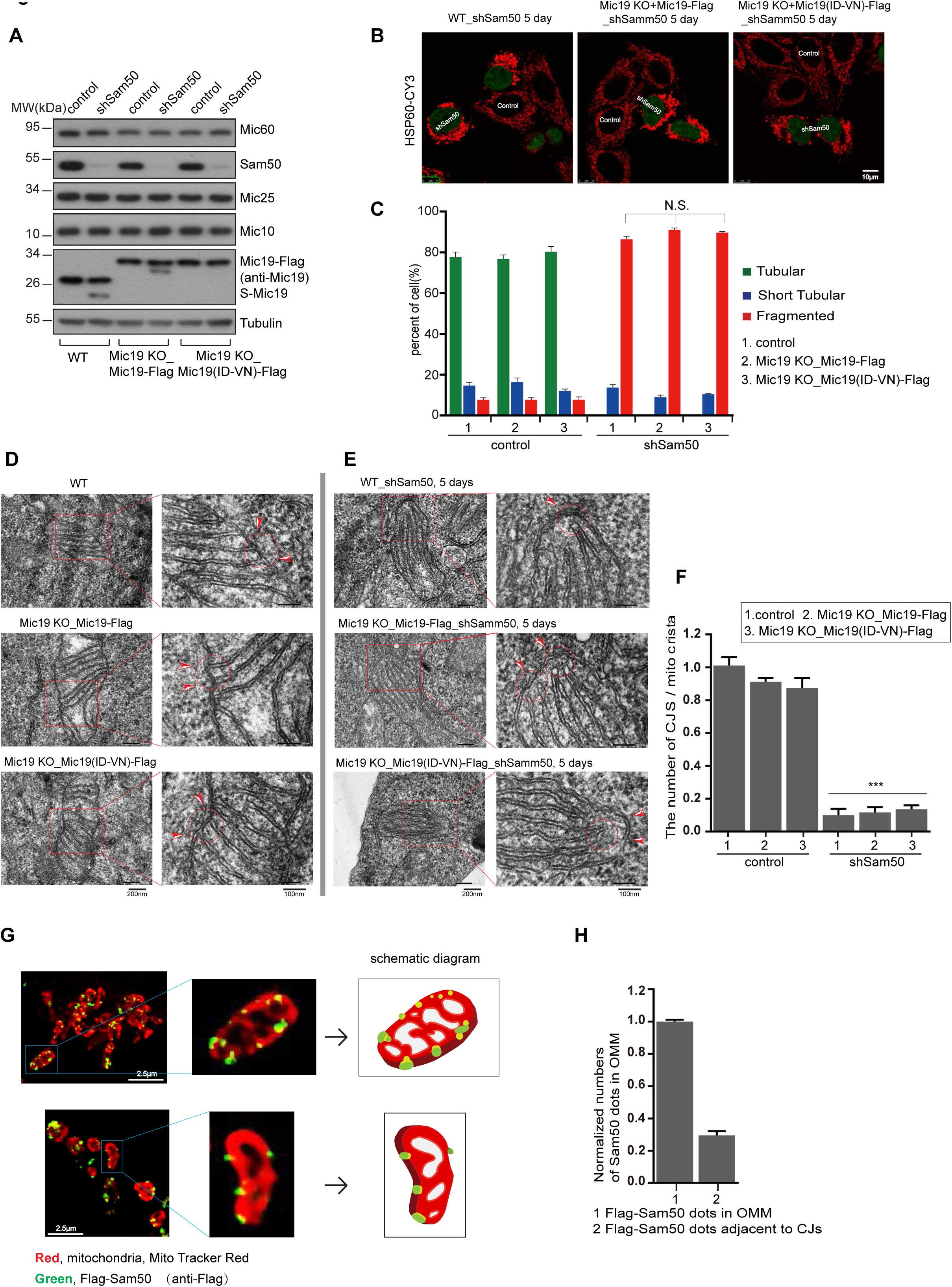
Sam50 may act as an anchoring point for Sam50-Mic19-Mic60 axis to guide the formation of mitochondrial crista junctions. (A) Knockdown of Sam50 (shSam50, 5 days) in WT, Mic19 KO, and Mic19^ΔgRNA^-Flag or Mic19 ^(ID^33-34^VN)ΔgRNA^-Flag expressed Mic19 KO HeLa cells were performed by infecting cells with control or shSam50 lentiviral particles, cell lysates were then analyzed by Western blotting using indicated antibodies. Tubulin was used as loading control. (B) WT, Mic19^ΔgRNA^-Flag, or Mic19 ^(ID^33-34^VN)ΔgRNA^-Flag expressed Mic19 KO HeLa cells were infected with control or shSam50 lentiviral particles. 5 days after infection, cells were fixed and immunostained with anti-HSP60 antibody, mitochondrial morphology was visualized by confocal microscope. (C) Mitochondrial morphology of indicated cell lines was classified into three types (tubular, short tubular, and fragmented) and counted. Bars represent means ±S.D. of three independent experiments. (D-E) The mitochondrial cristae junction structures of WT, Mic19^ΔgRNA^-Flag or Mic19 ^(ID^33-34^VN)ΔgRNA^-Flag expressed Mic19 KO HeLa cells with or without Sam50 knockdown were analyzed by transmission electron microscope (TEM). (F) Statistics of the mitochondrial ultrastructure in the indicated cell lines. The mean value and standard deviations (S.D.) were calculated from 3 independent experiments in which 100 mitochondrial cristae and the number of corresponding CJs were counted, ***P<0.001. (G) Flag-Sam50 stably expressed COS7 cells were stained with MitoTracker(r) Red CMXRos dye for labeling mitochondria, and then fixed and immunostained with Flag antibody, and visualized by stimulated emission depletion (STED) microscopy. The schematic diagram for Flag-Sam50 localization is showed. (H) Statistical analysis of Flag-Sam50 dots adjacent to the CJs structure.

### Restoration of Sam50-X-Mic60 axis by overexpression of Mic25 in Mic19 KO cells can reconstruct mitochondrial morphology and crista junctions

Following the results above, we have known that Sam50-Mic19-Mic60 axis plays a decisive role in the maintenance of mitochondrial morphology and structure, and destroying either of three subunits can disrupt the axis. However, large spherical mitochondria occurred in Mic60 knockdown cells ^17^ or Sam50 depletion cells ^36, 37^ but not Mic19 knockout cells (Figure 3A), suggesting that there may be another component (called X) which can partially replace Mic19 to form Sam50-X-Mic60 axis after Mic19 knockout. Among subunits of the MICOS complex, Mic25 may act as the candidate because Mic25 exhibits 36% overall sequence identity and 80% similarity to Mic19 ^13^. In addition, Mic60 depletion reduced Mic25, while Mic19 knockout decreased Mic60 but not altering Mic25 level (Figure S3A). Moreover, Mic25 is also capable of forming direct interaction with Sam50 and Mic60 (Figures S3B-S3D). Therefore, we investigate whether there is functional overlap between Mic19 and Mic25. Mic25 knockout did not affect the level of other MICOS subunits and mitochondrial morphology (Figures 5A and 5E). However, Mic25 knockdown (shMic25) in Mic19 KO cells caused mitochondria shape converting to ‘large spherical’ (Figures 5B and 5D), and led to an obvious further reduction of Mic60, Mic10 and Sam50 (Figure 5E). The similar phenotypes were observed in Mic19 knockdown (shMic19) plus Mic25 KO cells (Figures 5C and 5D). These results demonstrate that Mic25 functionally overlap with Mic19, but Mic19 play a dominant role in mediating Sam50-X-Mic60 axis and maintaining mitochondrial morphology.

**Figure 5.**
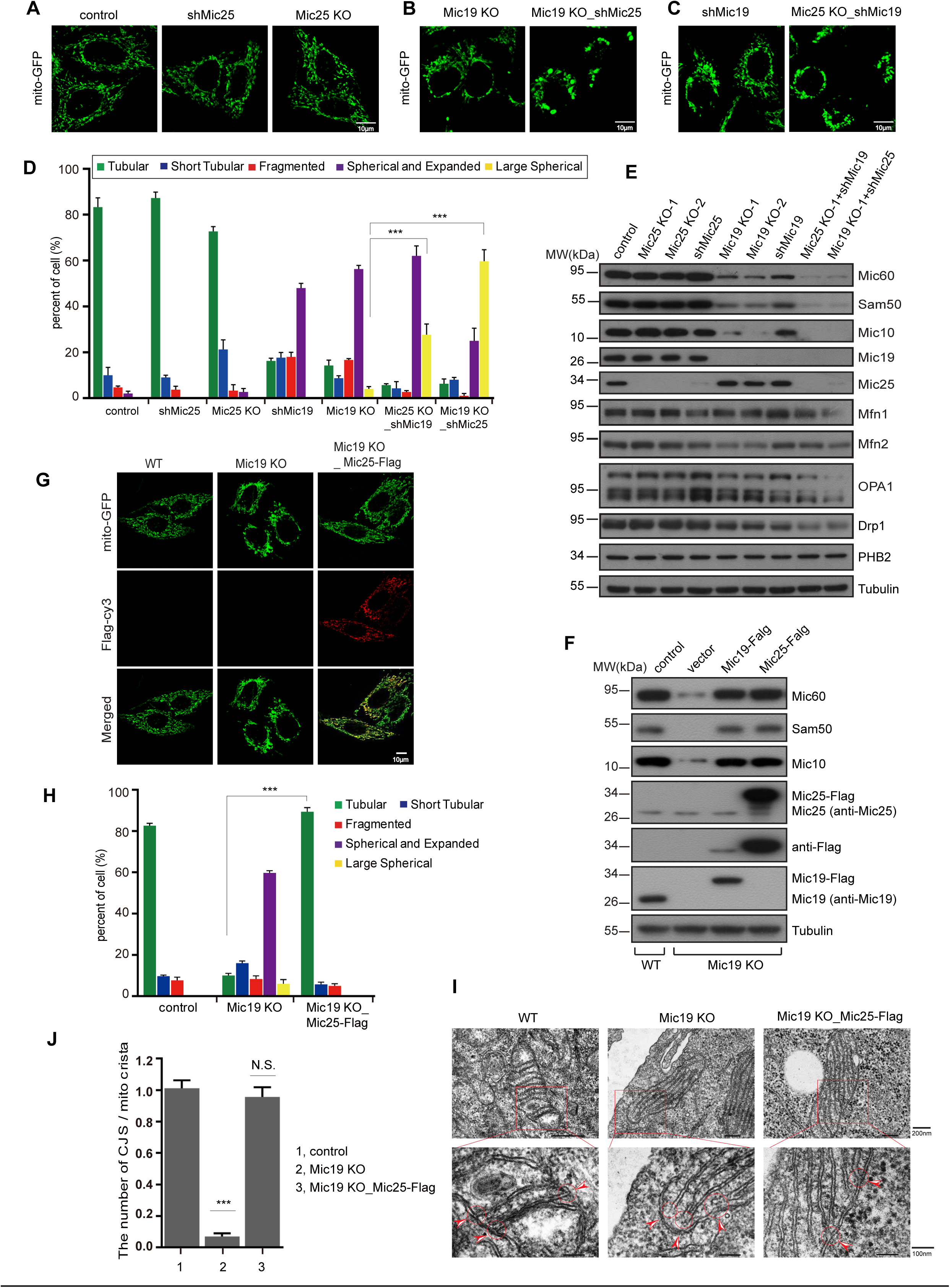
Restoration of Sam50-X-Mic60 axis by overexpression of Mic25 in Mic19 KO cells can reconstruct mitochondrial morphology and crista junctions. (A-C) WT (A), Mic19 KO (B), or Mic25 KO (C) HeLa cells expressing mito-GFP were knocked down for Mic19 (shMic19) or Mic25 (shMic25) respectively, and mitochondrial morphology was analyzed by confocal microscope. (D) Mitochondrial morphology in cell lines described in “A-C” was classified and counted according to the criteria detailed in “Materials and Methods”, and error bars represent means ±S.D. of three independent experiments, ***P<0.001. (E) Cell lysates of the indicated cells described in “A-C” were analyzed by Western blotting using the indicated antibodies. Tubulin protein was used as loading control. (F) Cell lysates of WT, Mic19 KO, and Mic19-Flag or Mic25-Flag overexpressed Mic19 KO HeLa cells were analyzed for protein levels by Western blotting using the indicated antibodies. (G) Mic25-Flag was stably overexpressed in Mic19 KO cells expressing mito-GFP, and immunostained with anti-Flag antibody. Mitochondrial morphology in WT, Mic19 KO, and Mic25-Flag overexpressed Mic19 KO HeLa cells was assessed by confocal microscope. (H) Statistical analysis of mitochondrial morphology in the cells described in “G” was performed according to the criteria detailed in “Materials and Methods”, the mean value and standard deviations (S.D.) were calculated from 3 independent experiments in which 100 cells were scored, ***P<0.001. (I) Mitochondrial crista junction structure of WT, Mic19 KO, or Mic25 overexpressed Mic19 KO HeLa cells were analyzed by transmission electron microscope (TEM). (J) Statistics of the mitochondrial ultrastructure in the indicated cells described in “H”. The mean value and standard deviations (S.D.) were calculated from 3 independent experiments in which 100 mitochondrial cristae and the number of corresponding CJs were counted, ***P<0.001.

Because Mic19 possess dominant role in Sam50-X-Mic60 axis comparing with Mic25, we then investigated whether Mic19 protein abundance is much higher than that of Mic25. We compared the relative protein levels of endogenous Mic19 and Mic25 in cells by comparing normalized Mic19-Flag with normalized Mic25-Flag. The level of endogenous Mic19 and Mic25 are comparable, and Mic25 slightly lower than Mic19 (Figures S3E-S3H), which further confirmed that the dominant role of Mic19 in Sam50-X-Mic60 axis. In addition, Mic25 KO cells display normal mitochondrial cristae architecture (Figures S3I and S3J). These data suggest that Sam50-Mic19-Mic60 axis plays the key role in mitochondrial cristae organization. Therefore, under normal conditions, Sam50-Mic19-Mic60 axis mainly exist; after Mic19 knockout, the unaffected Mic25 works together with the rest Mic60 and Sam50, and forms Sam50-Mic25-Mic60 axis to partially maintain the function of Sam50-X-Mic60 axis.

Since Mic25 can act as ‘spare bearing’, we investigated whether overexpressed Mic25 in Mic19 KO cells can form an axis to restore the mitochondrial outer and inner membrane contact. Overexpressed Mic25-Flag (about 10 fold of endogenous Mic25) in Mic19 KO cells recovered normal mitochondrial morphology (Figures 5G-5H). In addition, the level of other MICOS subunits was also recovered in Mic19 KO cells overexpressing Mic25-Flag (Figure 5F). These results suggested that overexpressed Mic25 could fill the position of Mic19 to form a new dominant axis with Sam50 and Mic60 and then reconnect the mitochondrial outer- and inner-membrane contact. Moreover, we also examined whether overexpression of Mic25 can reshape the ultrastructure of mitochondria. Interestingly, mitochondrial crista junctions reconstructed in Mic25 overexpressed Mic19 KO cells (Figures 5I and 5J). In addition, the reduced ATP level due to Mic19 deletion was recovered to the normal level when Mic25 was overexpressed in Mic19 KO cells (Figure S3K). Therefore, our data indicate that Mic25 is a compensatory factor for establishing Sam50-X-Mic60 axis, which bridges mitochondrial outer and inner membrane contact and regulates mitochondrial morphology and ultrastructure.

### The model of mitochondrial cristae organization

Based on previous reports and our findings, we formulate a theoretical model to illustrate mitochondrial cristae organization and the de novo formation of crista junctions (Figure 6):

**Figure 6.**
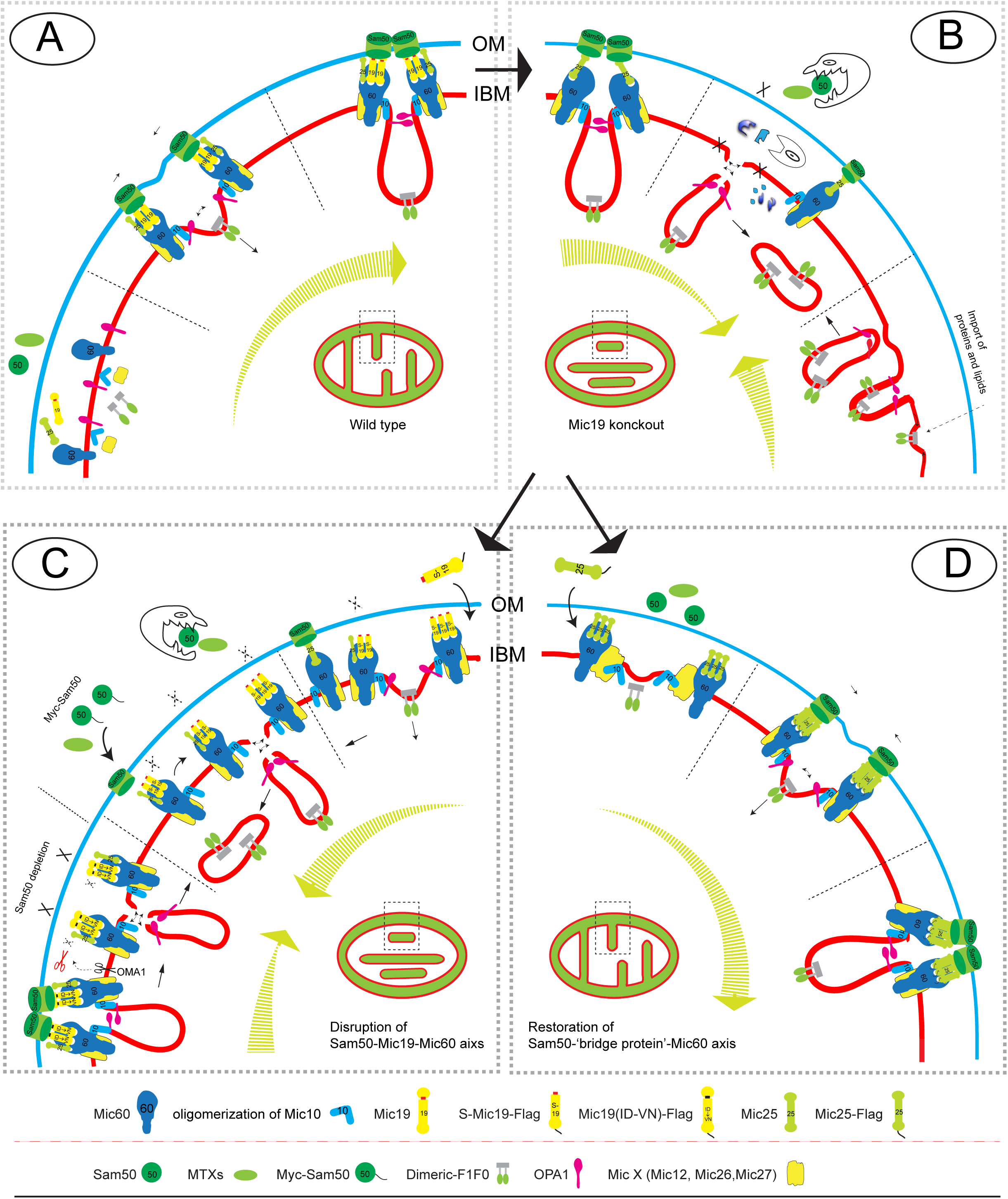
Theoretical model of Sam50-Mic19/Mic25-Mic60 axis functions on the formation of the CJs structure. In this schematic, the mitochondrial outer membranes and inner membrane form an arc. In the arc, we simulated the formation of mitochondrial cristae and CJ structures. Inside of the arc, the large arrow represents the direction of the formation or disappearance for mitochondrial cristae and CJ structures. If the tail of the large arrow changes from ‘thin’ to ‘thick’, it indicates the formation process. If the tail of the large arrow changes from ‘thick’ to ‘thin’, it indicates the disappearance process. In the innermost of the arc, simulated mitochondria represent the final mitochondrial cristae and CJs structure in each model 【 CJs, cristae junction site; Blue lines, the mitochondrial outer membrane (OM); Red lines, the mitochondrial inner boundary membrane (IBM) and cristae membrane (CM) 】.

A. When de novo cristae formed, monomeric F1FO assembled into dimeric F1FO providing positive curvature to crista membranes ^11, 14^. At the same time, Mic60 targets to inner membrane forming an anchor site, which is called nucleation, to recruit other MICOS subunits including Mic19, Mic10, and Mic25. Mic10 oligomerization can bend mitochondrial inner membranes and then stretch the mitochondrial inner membrane toward the matrix ^28^. Mic19 directly binding to Mic60, stretches out its N-terminal to physically interact with Sam50, and then close the distance between mitochondrial outer- and inner-membrane to form the contact site. OPA1 oligomerization can be used to regulate the contraction of the cristae junction (CJ) ^44^ ^45^,. In the end, curvature and fusion of inner membranes are halted when the integrated Sam50-Mic19-Mic60 axis is formed, meanwhile, CJ is generated.

B. upon Mic19 depletion, the Sam50-Mic19-Mic60 axis was disrupted, and then the associated subunits Sam50, MTXs, Mic60, and Mic10 were degraded by corresponding proteases, resulting in MIB complex was interrupted and depolymerized. Although there is still the Sam50-Mic25-Mic60 axis, it is not enough to maintain the contact between mitochondrial inner- and outer-membranes. Therefore, as there is no ‘blocking effect’ of the Sam50-Mic19-Mic60 axis, the CJs structure continues to shrink under the regulation of OPA1 and other factors and finally falls off from the inner membrane. At the same time, the nascent cristae membrane continues to produce in the absence of the Sam50-Mic19-Mic60 axis, and the membrane fusion proceeds as that in WT cells, but cannot halt ^14^. Thus, the final cristae forms are the stacked closed membrane sheets lacking connection with the IBM.

C. Expression of S-Mic19-Flag in Mic19 KO cells restores the MICOS complex, but without restoring Sam50, the Sam50-Mic19-Mic60 axis cannot be formed. Although, when S-Mic19-Flag and exogenous Myc-sam50 were simultaneously recovered, the Sam50-Mic19-Mic60 axis still was not established due to the defect of Mic19 N-terminal. Therefore, although all the elements, dimeric F1FO, OPA1, MICOS complex and SAM complex, are available, the CJ structure cannot be formed in the absence of interaction of Sam50-Mic19.

D. Mic25, which is not affected in the absence of Mic19, could form a Sam50-Mic25-Mic60 axis with the ‘residua’ Sam50 and Mic60. However, this axis is not sufficient to maintain CJ due to loss of the dominant Sam50-Mic19-Mic60 axis. When Mic25 is overexpressed, plentiful Mic25 will recruit more Sam50 and Mic60 to generate more Sam50-Mic25-Mic60 axis. When a sufficient number of axises work together, it will reconnect the contact between outer- and inner-membrane. As a result, the inner mitochondrial membrane structure CJ can be reconstructed.

## DISCUSSION

MICOS (mitochondrial contact site and cristae organizing system) complex is necessary for the formation of cristae organization and contact site between mitochondrial outer membrane and the inner boundary membrane. In the mammal, multiple subunits of MICOS complex were identified ^7, 38^. However, how the contact between mitochondrial outer and inner membrane and crista structure are formed remain elusive. In this study, we report that Sam50-Mic19-Mic60 axis serves as a dominant ‘bridge’ to connect mitochondrial outer and inner membrane; moreover, Mic19 actually acts as a dominant bearing to link SAM and MICOS complex and form a supercomplex MIB by Sam50-Mic19-Mic60 axis. Sam50-Mic19-Mic60 axis regulates mitochondrial morphology and determines mitochondrial cristae architecture.

We discovered that Mic19, the component of MICOS complex, can be cleaved at N-terminal by mitochondrial metalloprotease OMA1 when Mic19 detached from Sam50-Mic19-Mic60 axis either by depleting Sam50 or overexpressing Mic19. Why was little Mic19 cleaved under normal conditions (Figure 1C)? It is probably because the MICOS complex is assembled efficiently and precisely in proportion ^17^, the expression of Mic19 is strictly regulated and there is no ‘dissociative’ Mic19; additionally, Sam50 bound to its N-terminus of Mic19 may make the cut site to be hidden. Thus, Mic19 cannot be cleaved under normal condition. However, the stresses, which cause Sam50 downregulation, should induce Mic19 cleavage. Although the physiological relevance of Mic19 cleavage still unknown, this feature provides us with great tools to explore the key role of the Sam50-Mic19-Mic60 axis in combining mitochondrial outer and inner membrane contact. The cleavage of Mic19 interrupted the Sam50-Mic19-Mic60 interaction axis, and then resulted in the disruption of MIB complex. Although Mic60 can also interact with Sam50 ^33^, we have shown that Mic60 can be restored when expressed S-Mic19-Flag in Mic19 KO cells, but Sam50 is not restored (Figure 2F), suggesting that Mic60 alone cannot stabilize Sam50. In fact, Mic19 acts as the middle bearings to bridge and stable the outer membrane protein Sam50 and the inner membrane protein Mic60. It has been reported that N-terminal myristoylation of Mic19 may be required for the interaction with the outer membrane protein, and the myristoylation and the CHCH domain are essential for the import and mitochondrial localization of Mic19 ^32^, thus, S-Mic19-Flag losing its ability to combine with Sam50 may due to the missing of N-terminal myristoylation. However, S-Mic19-Flag still have the ability to import into mitochondria and assembly MICOS complex (Figures 2E and 3A).

In the last dozen years, MICOS and its subunits have been identified to be critical for mitochondrial membranes contact and cristae organization ^7, 25, 46^. However, Mic60 or Mic19 deletion always accompanies with the disruption of whole MICOS complex ^17, 18^. Therefore, it is hard to verify whether Mic60 or Mic19 alone is sufficient to regulate the cristae structure and mitochondrial membranes contact. Recently, Mic60 was reported to have the ability to bend liposomes in vitro ^25^, making Mic60 being a dominant player in stabilizing and promoting cristae formation. In addition, in vitro assay also revealed that Mic10 oligomerization could efficiently bend liposomes ^28^. These results suggest that Mic60 and Mic10 are cristae organization regulating effectors, and MICOS complex itself is supposed to be sufficient for cristae formation and organization. However, depletion of Mic60 or Mic19 not only severely repressed MICOS subunits but also mitochondrial outer membrane Sam50 (Figure S4A). In addition, Sam50 is also closely involved in cristae organization ^36, 37^. Therefore, it is questionable whether the abnormal cristae is caused directly by depletion of Mic60, Mic19 or Sam50. In our study, we proved that Sam50-Mic19-Mic60 axis is critical for MIB assembly and maintenance of normal mitochondrial morphology and ultrastructure. Disrupted Sam50-Mic19-Mic60 axis, even in the presence of SAM and MICOS complex, cause the abnormal mitochondrial morphology, the loss of cristae junctions, and the decrease of ATP production (Figure 3). Overall, Sam50-Mic19-Mic60 axis mediated MIB supercomplex, but not MICOS or SAM complex, determines mitochondrial cristae architecture.

The mechanism of formation and maintenance of mitochondrial crista junction (CJs) remains unclear, and most of the previous studies focus on the role of mitochondrial inner membrane proteins, such as OPA1, Mic60 and PHBs, in mitochondrial cristae organization, ^11^, 14, 45. However, it is rarely reported how mitochondrial outer membrane proteins act on the formation of CJs. We found that CJs fall off from the inner membrane when Sam50 was short-term depleted (5 days) without affecting MICOS core subunits Mic60 and Mic10 (Figure 4), indicating that outer membrane protein Sam50 is also indispensable for crista junctions formation. Additionally, Sam50-Mic19-Mic60 axis determines the mitochondrial cristae structure (Figures 3E and 3F), we put forward a new viewpoint that Sam50 acted as an anchoring point in mitochondrial outer membrane for the mitochondrial crista junctions. Therefore, Sam50 is not only functions in the biogenesis of mitochondrial β-barrel proteins, but also play role in the mitochondrial inner membrane organization. We also used STED to assess Sam50 localization, we expressed Sam50-Flag in HeLa cells for immunofluorescence assay by using anti-Flag due to the absence of available anti-Sam50 antibody for immunofluorescence. We observed about 30% Flag-Sam50 was detected over against to crista junctions (Figure 4H), further confirmed that Sam50 play a critical role in crista junction formation or maintenance.

Mic19 is highly conserved from yeast (Saccharomyces cerevisiae) to human. However, Mic25, an important paralog of Mic19, only exists in Metazoa ^25^, and the role of Mic25 in the MICOS complex is still unclear since Mic25 depletion does not affect mitochondrial ultrastructure (Figures 5A, S4I, and S4J). In this study, we revealed that there is a functional overlap between Mic19 and Mic25. Mic19 possesses dominant position to form Sam50-Mic19-Mic60 axis for mediating mitochondrial outer and inner membrane contact. While Mic25 function as a back-up component, which can substitute for Mic19 to form Sam50-Mic25-Mic60 axis with the residual Sam50 and Mic60 when Mic19 is depleted (Figures 5B-5E). Importantly, Mic19 could be cleaved by OMA1 under some stresses such as CCCP treatment, which results in disruption of Sam50-Mic19-Mic60 axis (Figures 3A-3D), but Mic25 could not be cleaved and remain stable (Figure 1H). Therefore, the Sam50-Mic25-Mic60 axis might protect the mitochondria to suffer some serious stresses. Mic19 plays a dominant role, whereas, Mic25 plays ‘Emergency Protective’ role when Mic19 was downregulated. It is likely not only to be a more economical, but also a more secure mechanism for the survival of multicellular organisms.

Taken together, Mic19 stabilizes and links Sam50 and Mic60 to establish Sam50-Mic19-Mic60 axis. The synergistic contact of mitochondrial outer and inner membrane mediated by Sam50-Mic19-Mic60 axis is critical for the establishment and maintenance of mitochondrial cristae architecture.

## ACKNOWLEDGMENTS

We gratefully thank Dr. Jiahuai Han (Xiamen University) for the gift of cDNA colony. Thank Dr. David Chan (California Institute of Technology) for communicating results before publication. This work is supported by National Natural Science Foundation of China (31471264 and 31671393), and the Fundamental Research Funds for the Central Universities (2042017kf0197 and 2042017kf0242).

## MATERIALS AND METHODS

### Cell culture and transfection

All cell lines (MEF, HeLa, HCT116 and 293T) were cultured in Dulbecco’s Modified Eagle Medium (DMEM) supplemented with 10% (v/v) fetal bovine serum (PAN, Germany), 100 U/ml Penicillin and 100μg/ml Streptomycin (Gibco) and 1mM L-glutamine at 37°C in 5% (v/v) CO2. Lipofectamine 2000 and Opti-MEM I (Invitrogen, Carlsbad, CA, USA) were used for transient transfection with expression constructs according to the manufacturer’s protocol.

### Antibodies and Reagents

Antibodies were used in this study: anti-Metaxin-2, anti-Prohibitin-2, anti-Mic60, anti-Yme1L and anti-SDHA were purchased from Proteintech; anti-Mfn1, anti-Mfn2 and anti-Sam50 were from Abcam; anti-Metaxin-1, anti-Mic19 and anti-HSP60 were from Abclonal; anti-OMA1 was from Santa Cruz Biotechnology; anti-Mic10 was from Origene; anti-Drp1 and anti-OPA1 were purchased from BD Biosciences. Reagents used in this paper were: Dimethyl sulfoxide (DMSO, Sigma-Aldrich); Carbonyl cyanide 3-chlorophenylhydrazone (CCCP, Sigma-Aldrich);

### Electron Microscopy

The procedure for transmission electron microscopy (TEM) was performed according to the previous report ^47^. The 100mM sodium cacodylate buffer was replaced by 100mM phosphate buffer without CaCl2. The sections were supported on copper grids and then post-stained in uranyl acetate for 10min and then in lead citrate for 15 min, and the stained sections were imaged onto negatives using a JEOL electron microscope operated at 80 kV (JEM-1400 plus, Tokyo, Japan).

### Measurement of ATP Production

Cellular ATP levels were measured using an ATP assay kit (Celltiter-Glo Luminescent Cell Viability Assay, Promega) according to the manufacturer’s instructions. Luminescence was measured using microplate reader and the values were normalized to the protein concentration.

### Statistical Methods

The data were presented as mean ± SD. Student’s t-test was used to calculate P values. Statistical significance is displayed as: N.S. (non-significance); *P <0.05, **P < 0.01, and ***P<0.001.

## SUPPLEMENTAL FIGURE LEGENDS

**Figure S1.**
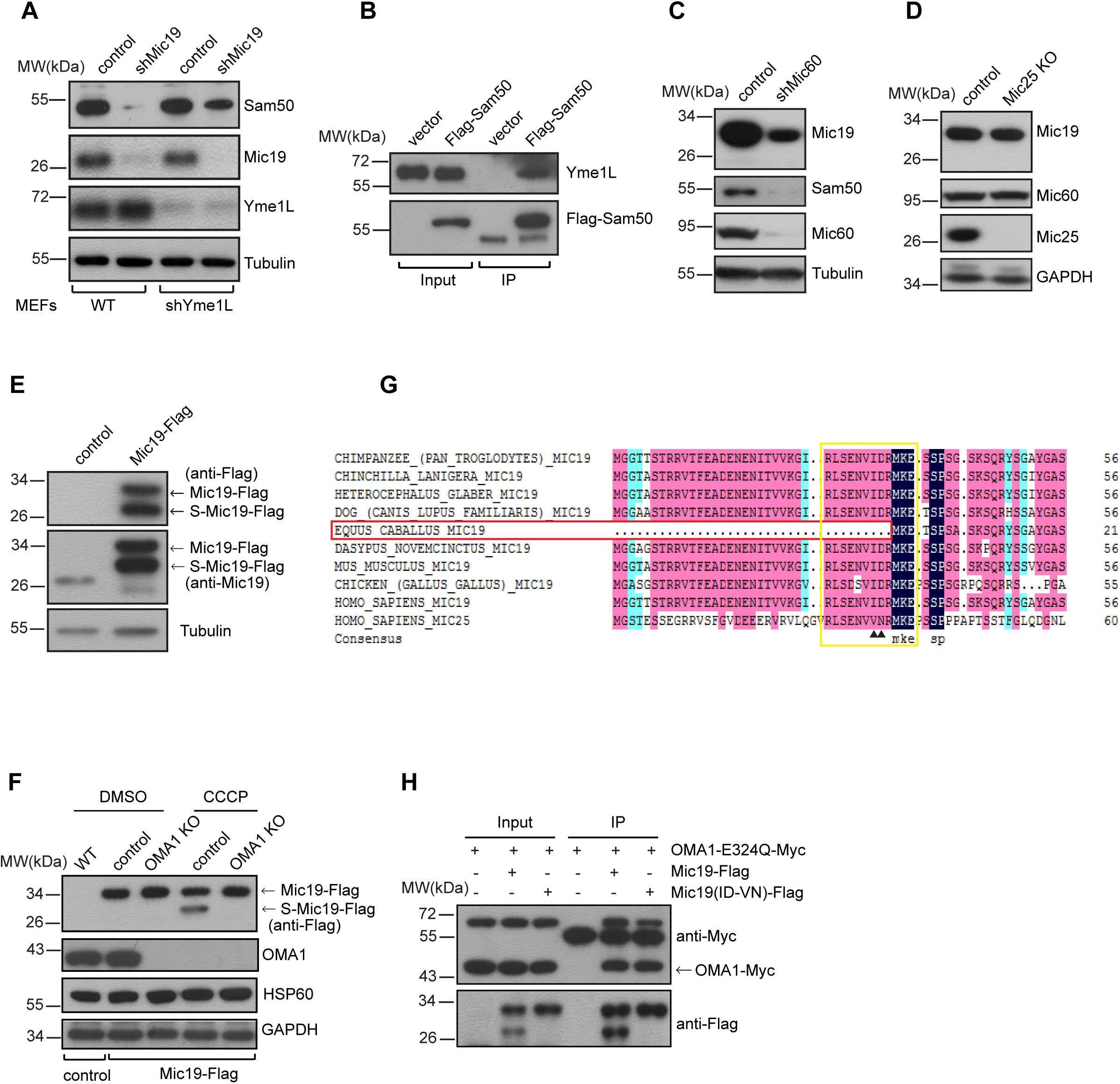
(Related to Figure 1) (A) Mic19 knockdown (shMic19) in WT or shYme1L (Yme1L knockdown) MEFs cells were performed respectively, and cell lysates were analyzed by Western blotting using the indicated antibodies. (B) Cell lysates of 293T cells expressing control or Flag-Sam50 were Immunoprecipitated with anti-Flag M2 resin, and the eluted protein samples were analyzed by Western blotting using anti-Yme1L or anti-Flag antibodies. (C-D) Cell lysates of control, Mic60 knockdown (shMic60, 5 days) MEFs cells (C), or Mic25 knockout HeLa cells were analyzed for Mic19 cleavage by Western blotting. (E) Cell lysates of control or Mic19-Flag transfected 293T cells were analyzed for Mic19 cleavage by Western blotting using anti-Mic19 or anti-Flag antibodies. Tubulin was used as loading control. (F) Mic19-Flag expressed WT or OMA1 KO HCT116 cells were treated with DMSO or CCCP (20μM, 4 h), cell lysates were analyzed for Mic19 cleavage by Western blotting. (G) The amino acid sequences of multiple species Mic19 and human Mic25 were obtained from NCBI, and then analyzed using sequence alignment software DNAMAN. (H) Cell lysates of 293T cells expressing Mic19-Flag plus OMA1^E324Q^-Myc or Mic19^ID^33-34^VN^-Flag plus OMA1^E324Q^-Myc were Immunoprecipitated with anti-Flag M2 resin, and the eluted protein samples were analyzed for interaction between Mic19 and OMA1 by Western blotting using anti-Myc or anti-Flag antibodies.

**Figure S2.**
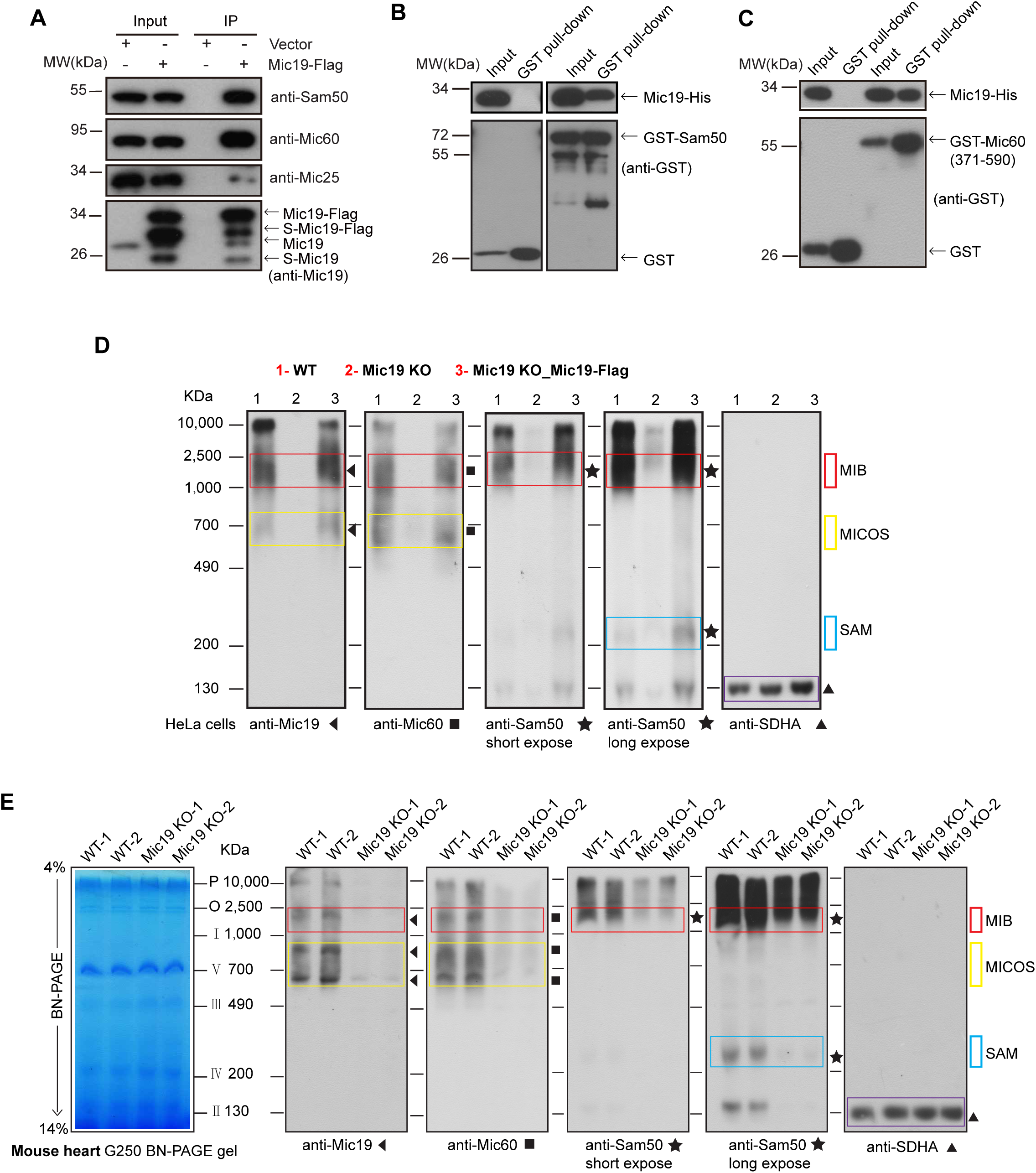
(Related to Figure 2). (A) Cell lysates of control or Mic19-Flag expressed 293T cells were used for co-immunoprecipitation (co-IP) assay with anti-Flag M2 resin. The eluted protein samples were detected by Western blotting analysis. (B-C) GST, GST-Sam50, GST-Mic60(371-590aa) or Mic19-His were expressed in E. coli, cell extracts were used for GST pull-down assay by using the Pierce Glutathione Agarose, and the protein samples were analyzed by Western blotting using anti-GST or anti-His antibodies. The examination of the direct interaction between Mic19 and Sam50 (B), or interaction between Mic19 and Mic60(371-590AA) (C). (D-E) Mitochondria were extracted from WT, Mic19 KO, or Mic19^ΔgRNA^-Flag expressed Mic19 KO HeLa cells (D), and from WT or Mic19 KO mice heart tissues (E). The non-denatured protein samples from mitochondria were analyzed for MIB, MICOS and SAM complexes by BN-PAGE and Western blotting using indicated antibodies. The bands of the complexes are labeled with corresponding boxes. Mitochondrial supercomplex II is detected by anti-SDHA, served as loading control.

**Figure S3.**
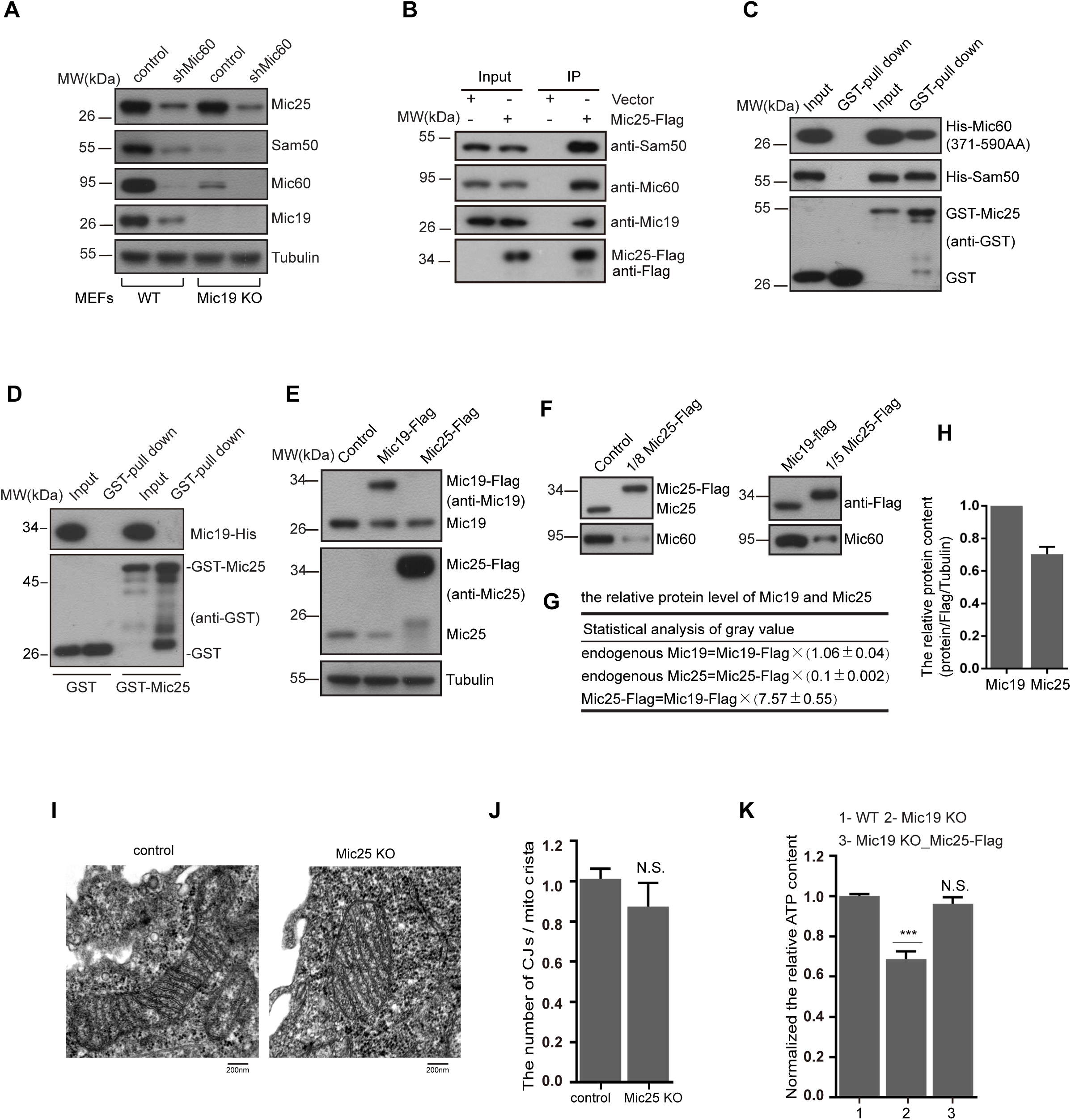
(Related to Figure 5). (A) WT or Mic19 KO MEFs cells were infected with lentiviral particles containing shMic60 for Mic60 knockdown, 5 days after infection, cells were lysed and used for Western blotting analysis using the indicated antibodies. (B) Lysates of control or Mic25-Flag expressed 293T cells were used for co-immunoprecipitation (co-IP) assay with anti-Flag M2 resin. The eluted protein samples were assessed by Western blotting analysis using the indicated antibodies. (C-D) GST, GST-Mic25, His-Mic60(371-590aa) or Mic19-His were expressed in E. coli, cell extracts were used for GST pull-down assay by using the Pierce Glutathione Agarose, and the protein samples were analyzed by Western blotting using anti-GST or anti-His antibodies. The examination of the direct interaction between Mic25 and Sam50 (C), the interaction between Mic25 and Mic60(371-590AA) (C), or interaction between Mic25 and Mic19 (D). (E-F) Lysates of HeLa WT cells expressing Mic19-Flag or Mic25-Flag (E) and the diluted protein lysates (F) were analyzed by Western blot analysis using the indicated antibodies. (G-H) Densitometry analysis was performed using ImageJ software to calculate the protein levels of related cells. The relative expression of Mic19 is set to 1, the expression of Mic25 was calculated according to the formulation in “G”, the data was shown in “H”. (I-J) The mitochondrial cristae junction structure of WT, Mic25 KO HeLa cells were analyzed by transmission electron microscope (TEM). The mean value and standard deviations (S.D.) were calculated from 3 independent experiments in which 100 mitochondrial cristae and the corresponding CJs were counted. (K) The relative ATP level of WT, Mic19 KO, or Mic25 overexpressed Mic19 KO HeLa cells were measured using an ATP assay kit. Error bars represent means ± SD of three independent experiments, ***P<0.001 vs WT.

## SUPPLEMENTAL EXPERIMENTAL TABLE

**Table S1.** Selected Tandem mass spectrometry (MS/MS) data from the eluted protein samples of co-IP assay.

| Gene Name | Description | MW<br>(kDa) | $\Sigma$#<br>Unique<br>Peptides | $\Sigma$# PSMs |
| --- | --- | --- | --- | --- |
| MIC60/IMMT | Mitochondrial inner membrane protein | 83.63 | 61 | 255 |
| MIC19/CHCHD3 | Coiled-coil-helix-coiled-coil-helix domain-containing protein 3 | 26.14 | 22 | 314 |
| MTX2 | Metaxin-2 | 29.74 | 13 | 38 |
| SLC25A5 | ADP/ATP translocase 2 | 32.87 | 5 | 18 |
| MIC25/CHCHD6 | Coiled-coil-helix-coiled-coil-helix domain-containing protein 6 | 26.44 | 8 | 12 |
| SAM50 | Sorting and assembly machinery component 50 homolog | 51.94 | 16 | 65 |
| HSP1A | Heat shock 70kDa protein 1A/1B | 70.01 | 20 | 64 |
| HSPD1 | 60kDa heat shock protein, mitochondrial | 61.02 | 10 | 18 |
| PGAM5 | Serine/threonine-protein phosphatase PGAM5 | 31.98 | 4 | 8 |
| ATP5C1 | ATP synthase subunit gamma, mitochondrial | 32.98 | 5 | 10 |
| ATP5B | ATP synthase subunit beta, mitochondrial | 56.52 | 7 | 13 |
| COX2 | Cytochrome c oxidase subunit 2 | 25.55 | 2 | 4 |
| UQCRC2 | Cytochrome b-c1 complex subunit 2, mitochondrial | 48.41 | 4 | 7 |

Table S1. Selected Tandem mass spectrometry (MS/MS) data. Mic19-Flag was transiently expressed in 293T cells, and co-immunoprecipitation (co-IP) assay was performed with anti-Flag M2 resin. The eluted protein samples were analyzed by MS/MS and the indicated proteins from MS/MS data was displayed in this table. The indicated protein of molecular weight (MW), the number and of total peptide spectrum matching (PSMs), the number of total unique peptides (Coverage) identified by MS are shown.

## SUPPLEMENTARY INFORMATION

### MATERIALS AND METHODS

#### Plasmids and shRNA constructions

The vector including pEGFP-N3, pMSCV-puro, pMSCV-hygro and Phage-puro were used to construct recombinant plasmids. shRNAi against target gene was performed using a modified retroviral vector with the H1 promoter to drive the expression of shRNAs ^1^. The following target sequences for gene knockdown were used: human Mic19: 5’-GGAGCTCAGAGTTCTACAG-3’; human Mic25: 5’-GACGCCGTGACACCTTCTA-3’; human Sam50-A: 5’-GGTCATCGATTCTCGGAAT-3’; human Sam50-B: 5’-ACATTCACTGAAATCATCT-3’; human Yme1L: 5’-GAGCTCTTCAAAGCATTTG-3’; human OMA1: 5’-GAAGTGCTTTGTCATCTAA-3’; Mouse Mic60: 5’-GGTGGTATCTCAGTATCAT-3’. Retrovirus production, cell infection and selection were performed according to the protocol described previously ^2^.

#### CRISPR/Cas9-mediated gene knockout

CRISPR/Cas9-mediated deletion of Mic19, Mic25, Mic10, Yme1L and OMA1, sgRNA sequences were selected using the MIT CRISPR design tool (http://crispr.mit.edu/) and cloned into LentiCRISPRv2 backbone (Addgene #52961). The following guide sequences were used: human Mic19: 5’- TCGGGAGAGGATATGTAGCG -3’; human Mic25: 5’-CTGATGCTGCCTTCGCCCGT -3’; human Mic10: 5’- TGTCTGAGTCGGAGCTCGGC -3’;human Yme1L: 5’- TGTCCAAGTGTTGGCCCCCG -3’; human OMA1: 5’-ACATTAGCATCCACCTCACG -3’. Lentiviral production, cell infection and selection were performed according to Zhang lab protocol ^3^.

#### Western blotting, Co-immunoprecipitation and GST-pull down assay

Western blotting and Co-immunoprecipitation assay were performed as described previously ^4^. For Western blotting, cell lysis buffer was 2×SDS sample buffer (0.125M Tris-Base, 4% SDS, 10% β-mercaptoethanol, 20% glycerol, 0.02% bromophenol blue, pH6.8). For Co-immunoprecipitation assay, cells were lysed with Triton X-100-based lysis buffer (1% Triton X-100, 10% glycerol, 20Mm Tris-Hcl, 150mM NaCl, 2mM EDTA, protease inhibitor mixture pH7.4).

For GST-pull down assay, GST, GST or His fused proteins in this study were expressed in coli. Plasmids coding for GST or His fused proteins were transformed cultured in E. coli BL21 at 37°C for 20-24h, then take 500μl bacteria solution to inoculate into 10ml LB liquid medium at 37°C for additional about 2∼3h. When the optical density reaches (OD600=0.6), the E. coli induced with 0.4mM∼1mM IPTG at 16°C for 18-24h. Cells were then harvested and suspended in 1× *E. coli* lysis buffer (1%Triton X-100, 150mM Tris-Hcl, 300mM NaCl, protease inhibitor mixture pH7.4) and disrupted by vigorous sonication on ice. Pierce Glutathione Agarose beads (Thermo Scientific, Waltham, MA, USA) were washed three times with ice-cold PBS, and mixed with supernatant containing GST and His fused proteins, the mixtures were rocked for 4h at 4 °C, followed by five washes of protein-bound beads with ice-cold PBS. Bound proteins were eluted and were analyzed by Western blotting.

#### Immunostaining

Cells grown on coverslips were fixed with 4% paraformaldehyde (dissolved in PBS PH7.4) for 15∼20min at room temperature and washed twice with PBS buffer. Then coverslips were incubated with 0.1% Triton X-100 (diluted with PBS PH7.4) for 10min and washed with PBS buffer, then blocked with 5% fetal bovine serum (FBS) in PBS for 1hr at room temperature. Next, cell coverslips were incubated with primary antibody (diluted with 5% FBS) for 1∼2hr at room temperature, then washed with PBS buffer three times per 5min, followed by secondary antibodies for 1hr at room temperature and washed with PBS buffer three times per 5min. Finally, cells were mounted and visualized by confocal microscopy.

#### Confocal Microscopy and Image Processing

Confocal images were collected using Leica microscope (Leica microsystem, Germany). All images for a given experiment were collected and adjusted in an identical manner. The Leica Application Suite software (Leica microsystem Corporation) was used for image processing and analysis. To determine mitochondrial morphology, cells were randomly selected for quantitative analysis and visually scored into five classifications (’Tubular’, ‘Short Tubular’, ‘Fragmented’, ‘Spherical and Expanded’ and ‘Large Spherical’).

#### Isolation of mitochondria and BN-PAGE analysis

Homogenization of cells or tissues and solubilization of mitochondria for BN-PAGE were performed according to the previous protocol ^5^. We optimized the procedure to isolate mitochondria of HeLa cells. A 100-mm dish of ∼90% confluent cells was digested and collected in a 2.0ml EP tubes, and the cell pellet re-suspended in cell homogenization buffer (83mM sucrose, 6.6mM imidazole/HCl, pH 7.0), then ultrasonic disrupt cell membranes on ice with the condition (1% Power, hit 1s, stop 4s, one minute in total). Centrifuge for 10 min at 700g to clear unbroken cells or cell nucleus, and the supernatant subsequently was centrifuged for 5 min at 10000g to collect the pellet containing the crude mitochondria. The pellet further re-suspended and centrifuge for 10 min at 10000g to collect the mitochondria. Isolated mitochondrial were re-suspended in solubilization buffer A (50mM sodium chloride, 50mM Imidazole/HCL, 2mM 6-aminohexanoic acid and 1mM EDTA PH7.0 at 4°C) on ice and lysed by the addition of ∼2% digitonin for 30min. The samples were separated on 4–14% blue native polyacrylamide gels. The gel was transferred to PVDF and immunoblotted with the anti-Mic60, anti-Mic19, anti-Sam50 and anti-SDHA antibodies respectively.

